# lncOriL, a novel polyadenylated mitochondrial lncRNA common to zebrafish and human

**DOI:** 10.64898/2026.03.26.714394

**Authors:** Tor Erik Jørgensen, Alex Wardale, Stanislava Wolf Profant, Cesilie Amundsen, Åse Emblem, Ingrid S. Joakimsen, Ole-Lars Brekke, Bård O. Karlsen, Igor Babiak, Steinar Daae Johansen

## Abstract

Even though teleost fish and mammals share the same mitochondrial gene content and organization, the teleost mitochondrial transcriptome is still poorly understood. We characterized the mitochondrial transcriptome during zebrafish (*Danio rerio*) early development by long-read direct RNA sequencing. All heavy-strand specific mRNAs were found to carry 3’ poly-A tails of approximately 50-60 residues, and the transcriptome profile was distinctive but practically invariant between stages. Three unusual transcripts were however noted. These included two mRNAs (COI and ND5 mRNAs), with significant 3’ untranslated regions corresponding to antisense gene sequences, and a previously not described noncoding RNA named here lncOriL. The ND5 mRNA was found to carry one third of all detected m^6^A methylation sites in the zebrafish mitochondrial transcriptome. The 313 nt-long lncOriL transcript had an abundance comparable to that of ND5 mRNA and it mapped to mitochondrial genome region covering the origin of light strand replication and four flanking antisense tRNAs. A mitochondrial tRNA-derived fragment (tiRNA5-Asn), with a 35 nt perfect pairing-potential to lncOriL, was present at all stages. Additional analyses including adult zebrafish, scissortail (*Rasbora rasbora*), and monkfish (*Lophius piscatorius*) strongly corroborate the results of COI mRNA, ND5 mRNA, and lncOriL transcript prevalence among teleost fish. Surprisingly, our findings in zebrafish were further supported by mitochondrial transcriptome analyses in domestic pig (*Sus scrofa*) and human (*Homo sapiens*), including tiRNA5-Asn commonly present in human tissues, suggesting that lncOriL is ubiquitously expressed and regulated in vertebrates.

**Author Summary:** Mitochondria contain their own genome and produce essential RNAs needed for energy production. Although fish and mammals share the same mitochondrial gene organization, less is known about how mitochondrial RNAs are processed and regulated in teleost. Using Nanopore direct RNA sequencing, we examined mitochondrial RNAs during early zebrafish development and discovered three unusual transcripts that include extended non-coding regions. Two of these molecules, COI and ND5 mRNAs, carry long 3′ untranslated regions formed by antisense gene sequences, suggesting previously unrecognized regulatory potential. We also identified lncOriL, a highly structured long noncoding RNA that spans the origin of light-strand replication and is abundant during development. Strikingly, the same RNA feature, including lncOriL and a matching tRNA-derived small RNA (tiRNA5-Asn), was found not only in zebrafish but also in human mitochondrial transcriptomes. These findings support conservation of regulatory mitochondrial RNAs across main groups of vertebrate species. Our work reveals a new layer of mitochondrial RNA regulation and expands the current understanding of how mitochondrial gene expression is controlled.

## Introduction

Teleost fish and mammals share a mitochondrial gene content, order and organization, which suggest a conserved mitochondrial gene expression pattern [1–3]. Vertebrate mitochondrial genomes (mitogenomes) are circular DNA molecules usually of only 16-17 kb in size coding for 37 conventional gene products. Among these are 13 protein coding genes (PCGs), all encoding hydrophobic inner mitochondrial membrane proteins important for oxidative phosphorylation [reviewed in 4,5].

The mitochondrial transcriptome in human cells has been investigated in more detail [6–8] and several of the human hallmarks appear conserved in teleost fish [2,9,10]. The 13 PCGs are transcribed and processed into 11 distinct mRNAs, 10 generated from the heavy (H)-strand promoter HSP2 and one (ND6 mRNA) from the promoter at the light (L)-strand. Only the H-strand specific mRNAs become polyadenylated. Two of the mRNAs are bicistronic (ND4/ND4L and A8/A6 mRNAs), while all the others are monocistronic. Interestingly, more than half of the UAA stop codons are made by posttranscriptional editing during mRNA polyadenylation.

Vertebrate mitogenomes also code for many noncoding RNAs (ncRNAs), some of them being taxa- or tissue-specific. Human long ncRNAs (mt-lncRNAs) are best studied; they include antisense to conventional mRNAs (lncCytB, lncND5, lncND6), various RNA chimers that involve LSU rRNA in sense or antisense configurations, and a variety of circular RNA species (recently reviewed by [11,12]). Only a few mt-lncRNAs have been found in teleost fish, and most of them map to the mitochondrial control region (CR) [2,9,13]. The biological roles of mt-lncRNAs are still obscure and functional evidence is in most cases lacking. However, a few rDNA and CR-derived mt-lncRNAs in human appear to be associated with cancer conditions [14,15].

The zebrafish (*Danio rerio*) is a popular model system in early developmental research due to advantages such as transparent embryos, short generation time, frequent spawning, and a well-established husbandry and laboratory maintenance protocols [16,17]. Among others, zebrafish has also become a well-established model system in studying mitochondrial diseases and neuropathology [18]. The mitogenome copy number decreases drastically during zebrafish early development, from more than 10^7^ copies per cell at 1-hour postfertilization to less than 100 at the 256-cell stage, an observation explained by lack of mitogenome replication during the early stages [19,20]. However, mitogenome transcription and corresponding RNA processing are active during all stages [19]. In this study, we investigated the mitochondrial transcriptome profile during zebrafish early development based on Oxford Nanopore Technology (ONT) direct RNA sequencing (dRNA-seq). Interesting findings were subsequently addressed in additional teleost fish species and in mammals, including human.

## Materials and Methods

### Zebrafish experiments, sample collection, and RNA preparation

Experiments were performed on zebrafish AB laboratory strain (ZIRC, University of Oregon, USA) at the zebrafish facility at the Nord University, Norway. All experiments and husbandry procedures were performed in agreement with the Norwegian Regulation on Animal Experimentation. Sample collection and RNA exraction from early developmental stages was performed and described previously [21,22]. The RNA samples for adult zebrafish were isolated and purified from skeletal muscle tissue and reported previously [10].

### RNA samples of scissortail, monkfish, and pig

The RNA samples for the scissortail (*Rasbora rasbora* S1-NU – also known as Gangetic scissortail rasbora) and monkfish (*Lophius piscatorius* BF2 – also known as European anglerfish) were isolated and purified previously from skeletal muscle tissue and heart muscle tissue, respectively, and reported in published studies [9,10]. The RNA sample from the pig (Norwegian landrace swine, *Sus scrofa domesticus*) was collected from liver and heart muscle tissue *post-mortem*, and total RNA was extracted. The pig was utilized in another study setting approved by The Norwegian Animal Research Authority, ID 31031.

### Human mitochondrial RNA

The human RNA samples were generated by using the human whole blood model with and without bacterial exposure, as described previously [23,24]. In two separate experiments, six healthy volunteers donated fresh venous whole blood which was either exposed to the gram-positive bacteria *Staphylococcus aureus* (Experiment 1) or the gram-negative bacteria *Escherichia coli* (Experiment 2) for 120 min (T120) at 37°C. Whole blood with phosphate-buffered saline (PBS) instead of bacteria was used as unstimulated control and did not have an incubation step (T0). At the end of the incubation, samples of whole blood with either PBS or bacteria were transferred to tubes containing PAXgene-solution and stored at -80°C until extraction. Total RNA, including mitochondrial RNA, was isolated using the MagMAX™ for Stabilized Blood Tubes RNA Isolation Kit (Ambion, Life Technologies) according to the manufacturer’s protocol. After RNA quality checks, equal amount of total RNA from the six donors with the same treatment where pooled together. Different blood donors were included in Experiments 1 and 2. The study was approved by the regional ethics committee of the Northern Norway Regional Health Authority. Bacterial strains: Experiment 1, *Staphylococcus aureus* ATCC 12598, final concentration 10^8^ cells/ ml. Experiment 2, *Escherichia coli* ATCC 25922, final concentration 10^7^ cells/ ml.

### ONT direct RNA sequencing

RNA-sequencing libraries were prepared following the Direct RNA Sequencing Kit (Oxford Nanopore Technologies, UK), namely SQK-RNA002 kit for *D. rerio* samples and SQK-RNA004 kit for *H. sapiens*, S*. scrofa*, *R. rasbora* and *L. piscatorius* samples. The optional RT step was used. The sequencing was performed on a MinION (Mk1B) equipped with FLO-MIN106 and FLO-MIN004RA flow cells, compatible with SQK-RNA002 and SQK-RNA004 kits, respectively.

### Mitochondrial transcriptome profile and poly-A tail assessment

Raw signal data (pod5 files) were base-called using Dorado v0.9.6 (Oxford Nanopore Technologies, UK) for the SQK-RNA002 kit, and Dorado v1.0.0 for the SQK-RNA004 kit. We employed the ‘*rna002_70bps_hac@v3*’ and ‘*rna004_130bps_sup@v5.2.0*’ base calling models for the SQK-RNA002 and SQK-RNA004 kits, respectively. The Dorado option ‘*--estimate-poly-a’* was used for poly(A) tail length estimation. Following basecalling, the raw BAM files were converted to FASTQ format using samtools (v1.22) [25]. The *‘samtools fastq -T ‘*’’* was used to preserve the ‘*pt’* tag (poly(A) tail length) information during the conversion. The reads were then aligned to their respective species-specific mitochondrial reference genomes using minimap2 (v2.29) [26] with parameter ‘*-ax splice -uf -k14’*.

Downstream analysis, quantification and visualization were performed in R (v4.5) (https://www.R-project.org). Mitochondrial transcript data were processed using the *‘Rsamtools’*, *‘GenomicAlignments’*, and *‘GenomicRanges’* libraries. Poly(A) tail lengths were extracted from the ‘*pt’* tag for defined mitochondrial regions, such as lncOriL (e.g., coordinates 5159–5476 for *D. rerio*). All non-numerical (NA) values and zero-length estimates were excluded from the analysis. The processed BAM files were imported to quantify mapping coverage across the mitochondrial genome and visualized alongside poly(A) tail length distribution. NanoCount (v1.0) [27] was employed for estimating transcript abundance and data was visualized in R (v4.5), using *‘dplyr’*, *‘tidyr’*, and *‘ggplot2’* libraries.

RT-PCR and Sanger sequencing were used as an alternative approach to map the poly-A sites of COI mRNA, ND5 mRNA, and lncOriL in adult zebrafish. The analysis was performed essentially as described previously [2], but with some minor modifications. First strand synthesis was generated by including the poly-A specific primer R-polyT (OP41: 5’-CGA CGC ATG CAC GCA TTT TTT TTT TTT TTT-3’) and the highly processive Induro reverse transcriptase [28]. Subsequently, the PCR reaction was performed using the tag-specific primer R-tag (OP313: 5’-AAG CG ACGC ATG CAC GCA TT-3’) and the upstream forward primer F-gene. R-tag binds to a unique sequence included in R-polyT. The F-gene primers used to generate amplicons from COI mRNA, ND5 mRNA, and lncOriL were ZM_6945_co1F (5’-AGT TGA ATT GAC CGC AAC AA-3’), ZM_13733_nd5F (GTC TTC CTT ACC TCT ATT AT-3’), and ZM_5178_aolF (5’-TAG GAC TTA CAG ACG TTA CT-3’), respectively. The amplicons were separated on an agarose gel, eluted, and Sanger sequenced as described previously [2].

### Mining tRNA-derived fragments from human database

Small noncoding RNA tissue atlas from 183 human samples was obtained from the Saaland cohort (PRJNA667263) miRNATissueAtlas 2025 [29]. Web link to the cohort can be found at https://www.ccb.uni-saarland.de/tissueatlas2025. Mined reads underwent adapter trimming and quality filtering (fastp v0.20.1, Q20, 15–30 nt), and duplicates were collapsed to unique reads with counts. RPM normalization accounted for library size differences.

### Mitochondrial RNA m^6^A site calling

Mapping N^6^-methyladenosine (m^6^A) modifications in mitochondrial mRNA was performed from our publicly available dataset [30] based on the m^6^A-SAC-seq approach [31,32]. Stringent m^6^A site calling only including the DRACH (D=G, A or U; R=A or G; H=A, C or U) consensus motif was performed as previously described [30]. Here, high confidence m^6^A sites were only called if they were found identical in at least two of the three replicates.

## Results

### Mitochondrial poly-A RNA expression during zebrafish early development

The mitochondrial mRNA expression was investigated during zebrafish early development at 1-cell, 32-cell, 256-cell, oblong, 50% epiboly, and bud stages. RNA isolated from adult muscle tissue was also included as a reference. The poly-A total RNA fraction was captured from each stage and subjected to library preparation and ONT dRNA-seq.

We noted several characteristic features when inspecting and comparing the mitochondrial mRNA profiles (Figure 1; S1 Table; S1 Figure). The same set of ten H-strand specific polyadenylated mRNAs was found in all developmental stages. The mRNAs were in general monocistronic, except for ND4L/4 mRNA and ATPase8/6 mRNA which were bicistronic. Complex I (ND) mRNAs were clearly less abundant compared to Complex IV (CO) mRNAs. The L-strand specific ND6 mRNA was, as expected, not detected due to lack of poly-A tail in all vertebrates.

**Figure 1.**
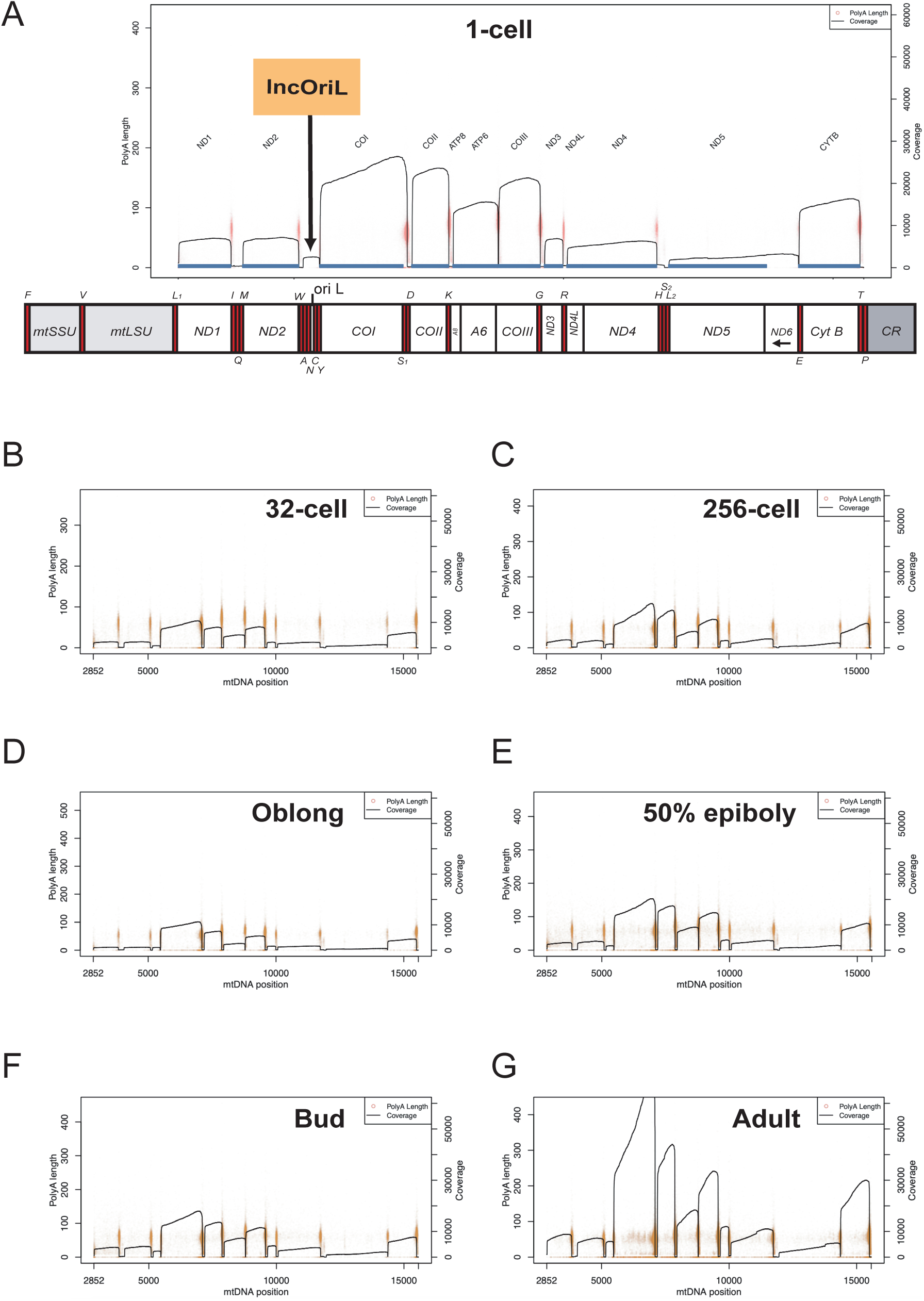
Mitochondrial transcriptome profiles in zebrafish during early developmental, based on ONT dRNA-seq. **A**) 1-cell stage. Poly-A length of individual RNA reads is indicated by red dots. Location of lncOriL is indicated. A linear map of the circular zebrafish mitogenome is presented below the profile. All mitochondrial genes are coded by the H-strand, except the L-strand specific ND6 and 8 tRNA genes. Gene abbreviations: mtSSU and mtLSU, mitochondrial small and large ribosomal DNA; ND1 to 6, Complex I NADH dehydrogenase subunit 1 to 6; CytB, Complex III cytochrome b; COI to III, Complex IV cytochrome oxidase subunit I to III; A6 and A8. Complex V ATPase subunit 6 and 8. CR, control region. **B-F**) Mitochondrial transcriptome profiles from stage 32-cell to Bud. **G**) Mitochondrial transcriptome profile from adult muscle tissue.

All PCG stop codons in the zebrafish H-specific mRNAs were UAA, and 7 of these were generated by post-transcriptional polyadenylation. Eight mRNAs had polyadenylation sites at or close to the UAA stop codon. However, COI and ND5 mRNAs carried longer 3’ untranslated region (3’ UTR) of 73 nt and 590 nt, respectively. Finally, a 313 nt-long polyadenylated ncRNA was detected between the ND2 and COI genes, covering the origin of light strand replication (OriL) and four flanking antisense tRNA genes. This previously not described ncRNA was named lncOriL.

### m^6^A modifications in mitochondrial RNA during zebrafish early development

An Illumina-based and publicly available m^6^A sequence dataset [30] that covers several zebrafish early developmental stages (1-cell, 256-cell, oblong, 50% epiboly, and bud) was used to extract high-confident m^6^A sites in mitochondrial mRNAs at nucleotide resolution. Only m^6^A sites embedded within the DRACH consensus motif were considered. We identified a total of 28 m^6^A sites in zebrafish mito-transcriptome that varied from only three in the oblong stage to twelve in the 1-cell stage (Figure 2; S2 Table). The ND5 mRNA was found to carry nine m^6^A sites. Three sites appeared fully methylated and located in ND5 mRNA and the anti-sense transcript to the CytB gene (lncCytB). Most m^6^A sites (94%) were detected at H-strand specific RNA, including Complex I (ND1, ND2, ND5), Complex III (CytB), Complex IV (COII, COIII), and Complex V (ATPase6, ATPase8) mRNAs. Together, we observed the mitochondrial m^6^A sites to be dynamic and reversible and with very few modified sites at the oblong stage, which corresponds roughly to the maternal-to-zygotic transition stage [30].

**Figure 2.**
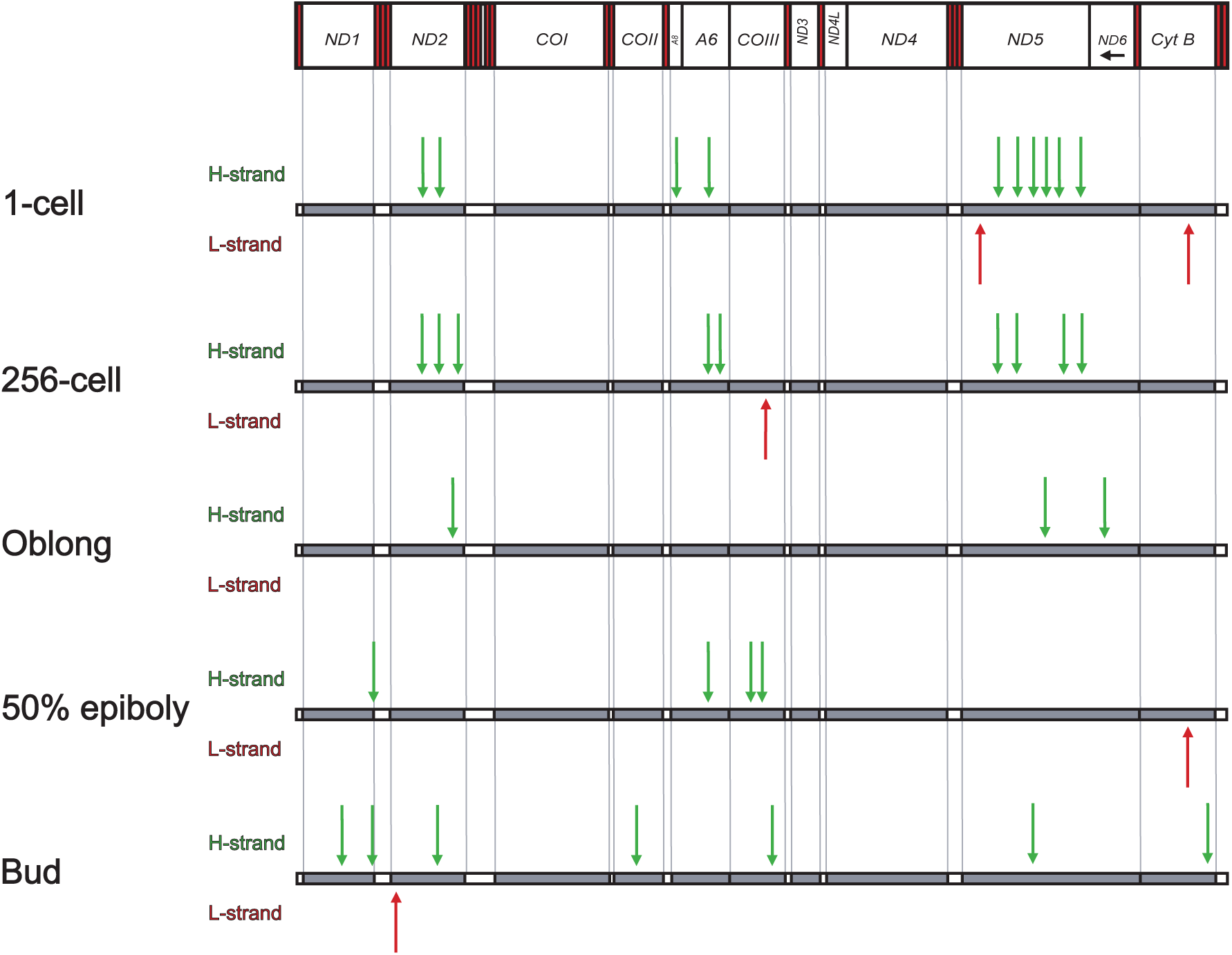
m^6^A mapping of zebrafish mitochondrial mRNAs during early development. Five embryonic stages were assessed: 1-cell, 256-cell, oblong. 50% epiboly, and bud. Green arrows, H-strand specific sites; red arrows, L-strand specific sites. See Supplementary Table S2 for additional information. Only DRACH-motives were included.

### COI and ND5 mRNA 3’ UTRs appear conserved in teleost fish and mammals

Transcriptome profile analyses revealed that the COI and ND5 mRNA carried 3’ UTRs corresponding to antisense mitochondrial genes. Two different experimental approaches were used to detect the 3’ UTRs in zebrafish. ONT direct RNA sequencing generated mainly full-length mRNA reads from the 3’ poly-A tail to the 5’ end (Figure 1; Figure 3). ONT sequencing revealed that the 3’ UTR in COI mRNA corresponded to antisense tRNA-Ser_1(UCN)_ and was capped by a poly-A tail of 53 adenosines (mean value; Figure 3A and S3 Table). Similarly, the ND5 mRNA carried a 3’ UTR corresponding to antisense ND6 and tRNA-Glu and a 53 adenosines (mean value) poly-A tail (Figure 3C; S3 Table). The second experimental approach was based on RT-PCR combined with Sanger sequencing. Here, the first-strand primer (R-polyT) binds to the poly-A tail but also carry a unique tag sequence (see Materials and Methods). Finally, the amplicon was generated by an upstream gene-specific forward primer (F-gene) and a downstream tag-specific reverse primer (R-tag). The results from analyses of COI mRNA and ND5 mRNA transcripts are shown in Figures 3A and 3C, respectively. The RT-PCR-Sanger approach revealed exactly the same results as for the ONT dRNA-seq.

**Figure 3.**
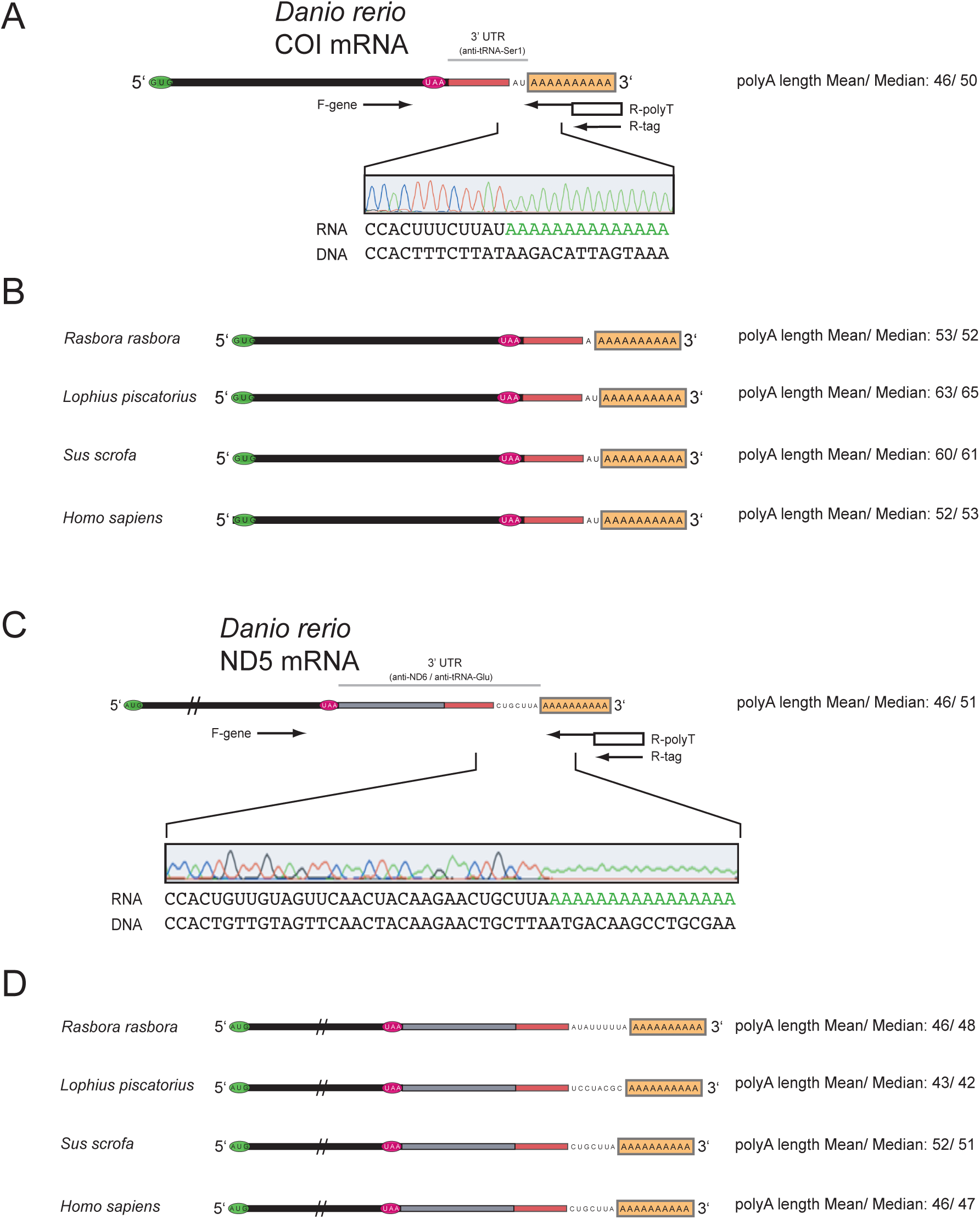
Mitochondrial 3’ UTR polyadenylation based on ONT dRNA-seq. **A**) Schematic presentation of the COI mRNA in zebrafish (*D. rerio*) carries a 73-nt 3’ UTR capped by a poly-A tail. The poly-A site was independently verified by a reverse transcriptase PCR-Sanger sequencing approach. R-polyT, first-strand synthesis primer that anneal to poly-A and carrying a unique tag-sequence. F-gene and R-tag, PCR primers that are gene-specific and tag-specific, respectively. The generated Sanger cDNA (RNA) sequence compared to the corresponding mtDNA sequence is shown below. **B**) Schematic presentation of the COI mRNA in scissortail (*R. rasbora*), monkfish (*L. piscatorius*), pig (*S. scrofa*), and human (*H. sapiens*). **C**) Schematic presentation of the ND5 mRNA in zebrafish (*D. rerio*) carries a 590-nt 3’ UTR capped by a poly-A tail. The poly-A site was independently verified by a reverse transcriptase PCR-Sanger sequencing approach. **D**) Schematic presentation of the ND5 mRNA in scissortail (*R. rasbora*), monkfish (*L. piscatorius*), pig (*S. scrofa*), and human (*H. sapiens*).

Next, we asked if the 3’ UTR features were unique to zebrafish. Total RNA was isolated and investigated in two additional teleost fish species:scissortail, a Danionidae species related to zebrafish, and monkfish, distantly related species, as well as in pig and human. The RNA samples were subjected to ONT dRNA-seq, and the poly-A RNA was subsequently mapped to their respective mitogenomes (S2 Figure). Our data showed that COI mRNA in all the investigated species carried the same 3’ UTR as determined in zebrafish and capped by a 3’ poly-A tail of approximately 50 adenosines (Figure 3B). Similar results were obtained from the ND5 mRNA analyses. In all the investigated species we found 3’ UTRs corresponding to a-ND6 and a-tRNA-Glu (Figure 3D).

### lncOriL, a polyadenylated mitochondrial lncRNA in vertebrates

In the mito-transcriptome profile (Figure 1) we identified a novel 312 nt polyadenylated ncRNA (lncOriL) mapping between ND2 and COI in the zebrafish mitogenome and being at an abundance comparable to ND5 mRNA (S1 Figure). lncOriL covered the antisense of four tRNAs (tRNA-Ala, tRNA-Asp, tRNA-Cys, and tRNA-Tyr) and the OriL hairpin (Figure 4A). A more detailed inspection of the ONT reads showed that the 5’ end of the zebrafish lncOriL corresponded exactly to the 3’ processing site of tRNA-W generated by RNase Z, and proposed to be highly structured (Figure 5A, left panel). The 3’ end of lncOriL was capped by a poly-A tail estimated to be 24% shorter than average poly-A of all mitochondrial transcripts (Figure 5A, right panel; S3 Table).

**Figure 4.**
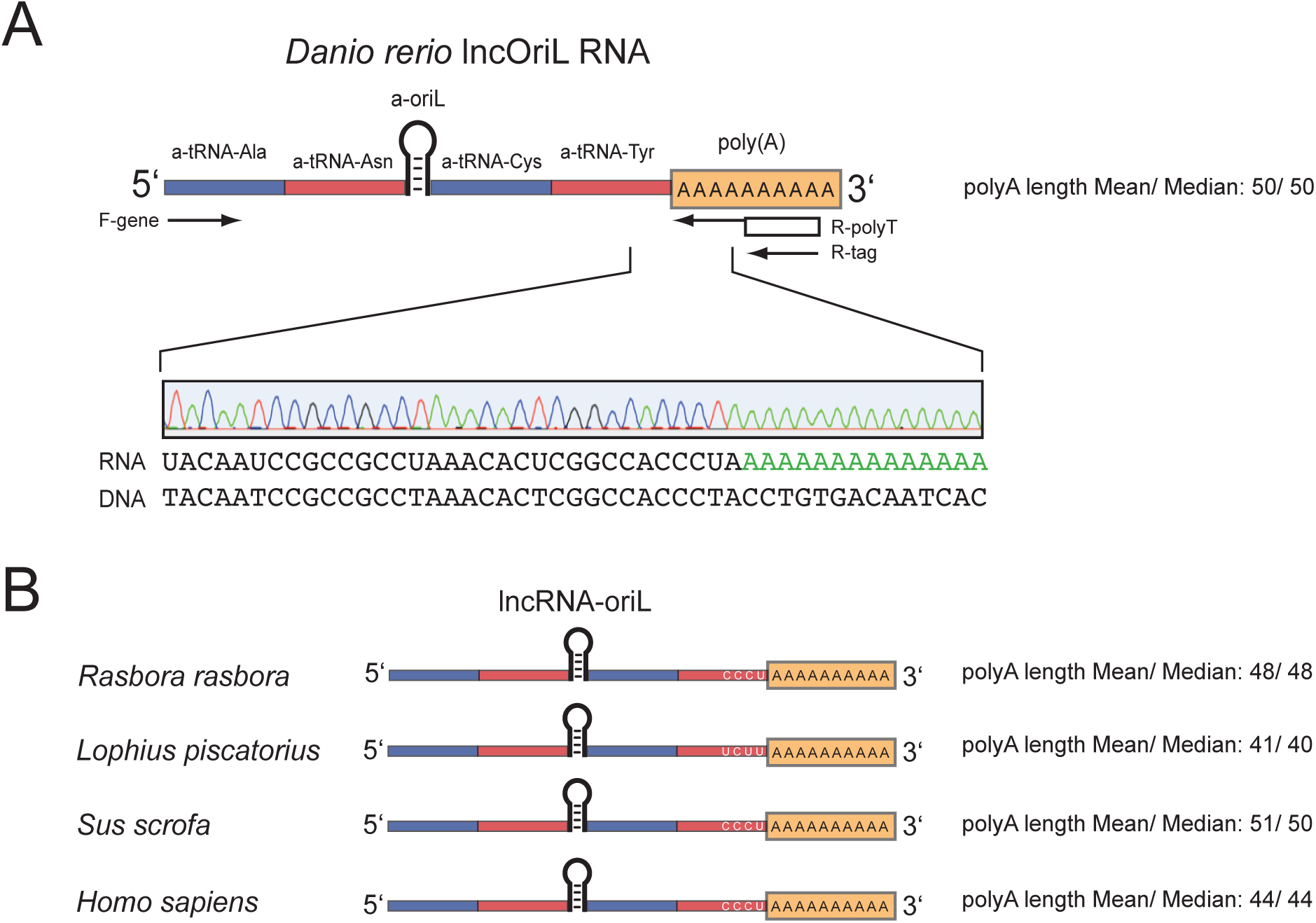
Key features of mitochondrial lncOriL. **A**) Schematic presentation of lncOriL RNA in zebrafish (*D. rerio*) based on ONT sequencing. lncOriL carries antisense oriL and tRNA sequences, as well as a poly-A tail of 50 residues (mean value). The poly-A site in lncOriL was independently verified by a reverse transcriptase PCR-Sanger sequencing approach. R-polyT, first-strand synthesis primer that anneal to poly-A and carrying a unique tag-sequence. F-gene and R-tag, PCR primers that are gene-specific and tag-specific, respectively. The generated Sanger cDNA (RNA) sequence is compared to the corresponding mtDNA sequence. **B**) Schematic presentation of the lncOriL RNA in scissortail (*R. rasbora*), monkfish (*L. piscatorius*), pig (*S. scrofa*), and human (*H. sapiens*).

**Figure 5.**
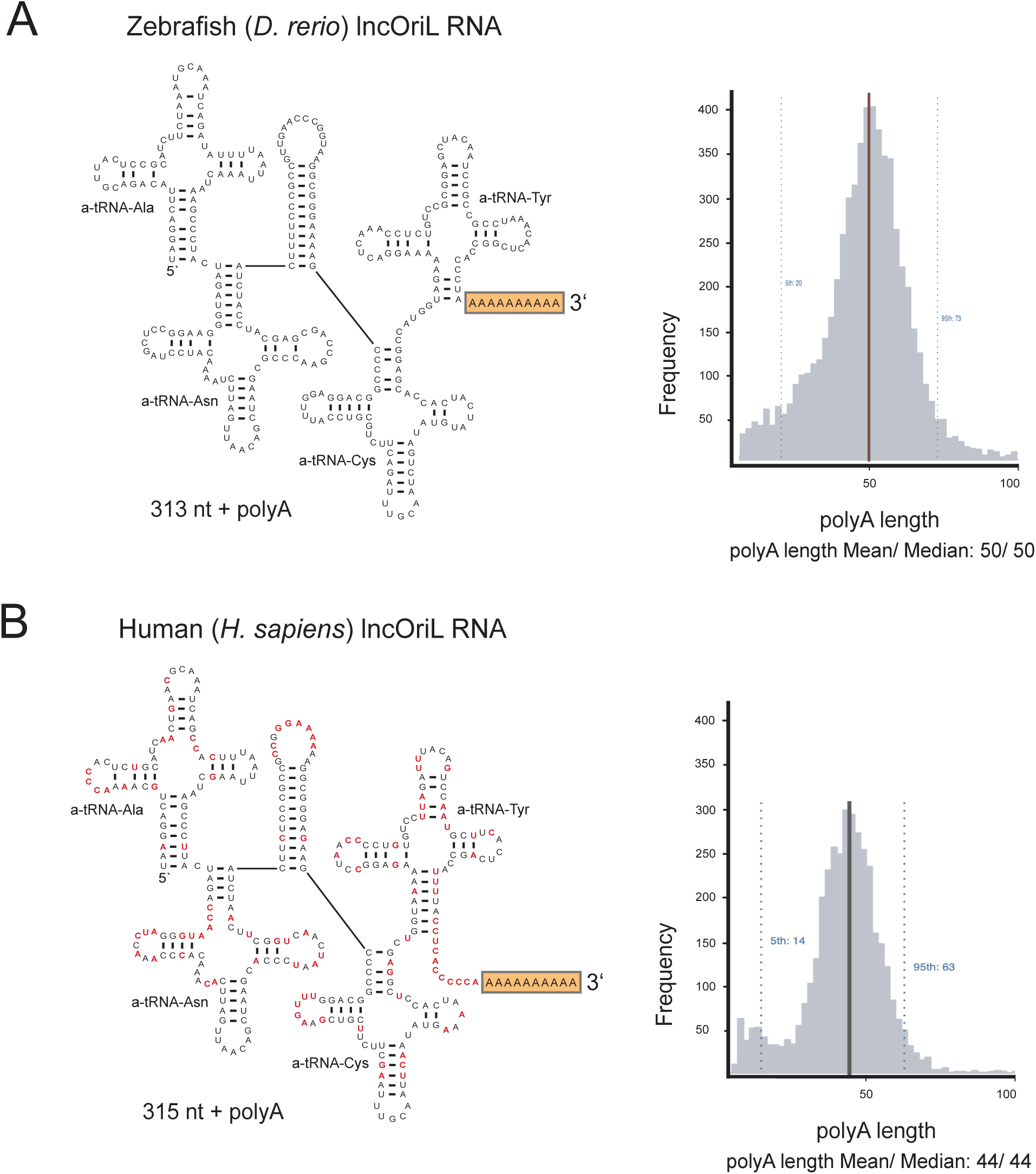
Secondary structure diagram of lncOriL. **A**) Zebrafish lncOril is 313 nt in size and has a poly-A tail of 50 residues. **B**) Human lncOril is 303 nt in size and has a poly-A tail of 44 residues and is 70% similar in nucleotide positions compared to that of zebrafish. Variable positions are indicated by red uppercase letters.

The poly-A feature of lncOriL RNA was verified by the RT-PCR-Sanger approach and found to be identical to that of ONT direct RNA sequencing (Figure 4A). We then asked whether lncOriL was present in other vertebrates as well. Indeed, lncOriL was detected in mito-transcriptomes of scissortail, monkfish, pig, and human (S2 Figure). Similarly to zebrafish, lncOriL transcripts had conserved organization, including the 5’ and 3’ ends, and a shorter than average poly-A tails compared to corresponding mRNAs (Figure 4B and S3 Table). Interestingly, despite the human lncOriL shared only 70% of nucleotide positions with zebrafish, the proposed RNA secondary structures in these two species were highly similar (Figure 5B).

### The mitochondrial lncOriL RNA is enriched in human blood cells

The human mito-transcriptome profile was assessed in cells from whole blood samples representing two different experiments, Ex1 and Ex2. In each experiment, a mixture of blood samples from six randomized human donors was exposed to bacterial stressors at 37°C. Ex1 and Ex2 were exposed to live *S. aureus* (10^8^ cells per ml final concentration) and *E. coli* (10^7^ cells per ml final concentration), respectively. RNA was isolated after 120 min exposure and directly sequenced by ONT. In parallel, the negative controls were included by replacing bacterial suspensions with PBS (phosphate-buffered saline). The mito-transcriptome profiles of control (T=0) and bacterial exposures (T=120) samples are shown in Figure 6. The overall profiles appear similar to those of zebrafish early development (Figure 1), and no clear differences were observed when comparing T=0 to T=120 in both Ex1 and Ex2.

**Figure 6.**
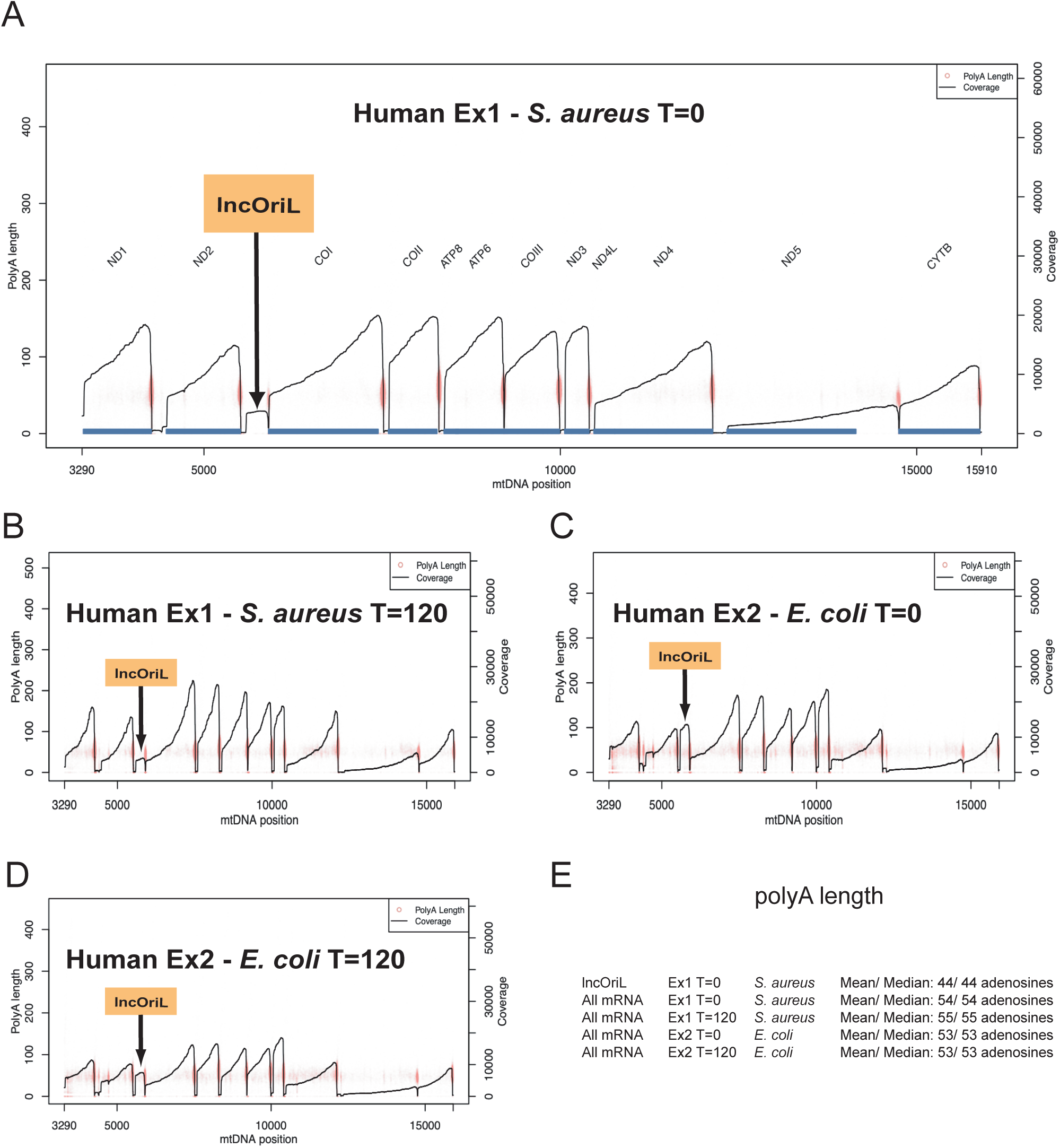
Mitochondrial transcriptome profiles based on ONT dRNA-seq of human blood samples exposed to *S. aureus* (experiment 1; Ex1) and *E. coli* (experiment 2; Ex2) bacteria. Ex1 and Ex2 contain a random mix of blood samples from six individuals. **A**) Ex1 transcriptome profile after 0 min, where *S. aureus* is replaced by PBS. lncOriL is indicated, and poly-A tail of individual RNA reads is shown by red dots. **B**) Ex1 transcriptome profile after 120 min exposure of *S. aureus*. **C**) Ex2 transcriptome profile at time 0 min transcriptome profile, where *E. coli* is replaced by PBS. **D**) Ex2 transcriptome profile after 120 min exposure of *E. coli*. **E**) The lncOriL poly-A length is approximately ten residues shorter than the transcription profile average.

However, we observed a distinct difference in the lncOriL enriched levels when comparing Ex1 and Ex2. While lncOriL in Ex1 was at an approximate same level as the ND5 mRNA, the same comparison in Ex2 revealed a 3 to 4 times relative increase in lncOriL (Figure 6A and C), consistent with individual variations in lncOriL expression level.

### The mitochondrial tRNA-derived fragment tiRNA5-Asn is antisense to lncOriL RNA

A subset of mtDNA-encoded tRNAs generates small RNAs with regulatory potential [33]. The mitochondrial tRNA-derived fragments in zebrafish were analysed in Illumina-generated and publicly available small RNA datasets from 1-cell, 32-cell, 128-cell, oblong, 50% epiboly, and bud stages. The small RNA datasets were mapped onto the zebrafish mitochondrial reference genome (PP963510). Each stage generated approximately 320,000 mapped mitogenome reads, corresponding to about 1% of the total mapped reads to the zebrafish genome. Six tRNAs (tRNA-Leu_1(UUR)_, tRNA-fMet, tRNA-Asn, tRNA-Lys, tRNA-Ser_2(AGY)_, and tRNA-Thr) were found to generate fragments corresponding to tRF5, tRNA 5’-half (tiRNA5), and tRNA 3’-half (tiRNA3) (S3 Figure). Two of the tRNA-derived fragments were ubiquitously detected. While tiRNA5-Ser_2(AGY)_ (30 nt) constituted the majority of the detected RNA fragments in zebrafish, tiRNA5-Asn (35 nt) was co-expressed with lncOriL at all developmental stages (Figure 7A). tiRNA5-Asn is of particular interest since it is antisense to lncOriL near the OriL hairpin structure (Figure 7B).

**Figure 7.**
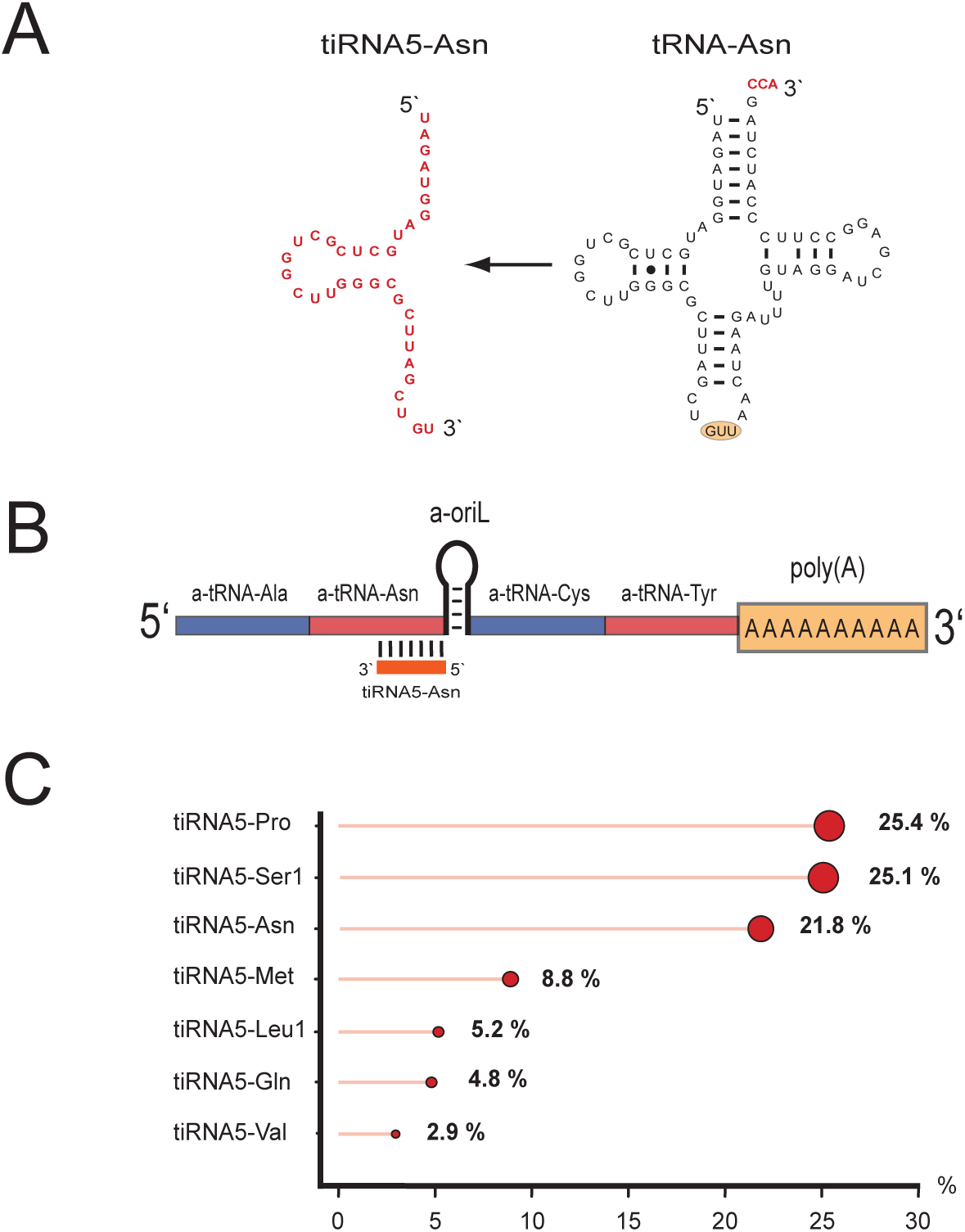
Mitochondrial tRNA-derived fragments in zebrafish and human. **A**) Mitochondrial tRNA-Asn in zebrafish generates the 35-nt tRNA-derived fragment tiRNA5-Asn in early development. **B**) tiRNA5-Asn makes a potential duplex (35 to 46 bp) in proximity to the OriL hairpin helix of lncOriL in zebrafish. **C**) Global abundance of mitochondrial tRNA-derived fragments in human tissue based on the Saarland cohort. tiRNA5-Asn accounts for 21.8% of all mitochondrial tRNA-derived fragments.

We then analysed 183 human samples from the Saaland cohort (PRJNA667263) spanning 61 distinct tissue types. Here, more than 8.6 million unique mitochondrial tRNA-derived fragment sequences were identified. The tissue types showed high individual integrity, and three tRNAs (tRNA-Pro, tRNA-Ser_1(UCN)_, and tRNA-Asn) accounted for the majority (72.3%) of all mitochondrial fragments detected (Figure 7C). Interestingly, they all represented the tiRNA5-type of tRNA-derived fragments. While tiRNA5- Ser_1(UCN)_ (40 to 46 nt) was antisense to the COI mRNA 3’ UTR, tiRNA5-Asn was antisense to lncOriL, varied in length from 38 to 46 nt, and possessed the same 5’ end as the zebrafish tiRNA5-Asn. Interestingly, tiRNA5-Asn was the only tRNA-derived fragment generated from an L-strand specific tRNAs in both zebrafish and human.

## Discussion

We have demonstrated that mitochondrial mRNA expression pattern is highly conserved between teleost fish and mammals, and that two of the ten polyadenylated mRNAs carry significant 3’ UTRs with an antisense regulatory potential. In addition, we identified an enriched but previously undetected polyadenylated noncoding RNA. This ncRNA, named lncOriL, covers the origin of light-strand replication and flanking antisense tRNA genes.

### Mitochondrial transcript profile and polyadenylation

All mature mitochondrial mRNAs, except the L-strand specific ND6 mRNA, are processed from the same polycistronic H-strand precursor transcript [34]. The relative mRNA expression profiles, however, show a characteristic pattern of enriched Complex IV mRNAs (COI-III mRNAs) compared to Complex I mRNAs (ND1 to 5 mRNAs). The overall profile was invariant during zebrafish early development, as well as similar among zebrafish adult tissue, scissortail, monkfish, pig and human. This characteristic profile appears linked to mRNA stability and supported by a recent report by Moran and co-workers [8] showing that Complex I mRNAs in human are significantly less structured compared to Complex IV mRNAs. Furthermore, the m^6^A modification in mRNA is also associated with mRNA stability [35], and our finding that 63 % of mitochondrial m^6^A sites are located at Complex I mRNAs (compared to only 17 % at Complex IV mRNAs) is in line with this notion. Polyadenylation is an essential maturation step in nuclear-generated eukaryote mRNA, and contributes to mRNA stability, translocation to cytoplasm, and translation at ribosomes [36]. However, roles and mechanisms of mitochondrial poly-A tails are still poorly understood [37]. We found all mitochondrial poly-A tails in mRNA to be at a similar length, ranging from 40 to 60 adenosine mean number, that included all zebrafish early developmental stages and adult tissues of the investigated teleost and mammals. This corroborates previous studies in human mitochondria [37].

### The 3’ UTR of COI mRNA

COI mRNAs in our dataset (zebrafish, scissortail, monkfish, pig and human) were found to be polyadenylated ∼75 nt downstream the UAA stop codon, generating a 3’ UTR corresponding to antisense tRNA-Ser_1(UCN)_. This is in line with previous findings in various mammals [38,39] and teleost fish [2,9,10]. No clear function has been assigned to the COI mRNA 3’ UTR, but one plausible explanation could be in control and regulation of RNA stability. The COI mRNA appears to be the most structured among the human mitochondrial mRNAs [8], an observation that corroborates our findings that this mitochondrial mRNA is the most enriched messenger across all species investigated. Interestingly, the Saarland cohort profiling identified tiRNA5-Ser_1(UCN)_ as one of the most expressed tRNA-derived fragments in humans, a small RNA (∼40 nt) that is complementary to the 3’ UTR in COI mRNA. Furthermore, studies in rat have shown that miR-181c, a nuclear encoded microRNA, specifically targeted the 3’ UTR in COI mRNA [38]. The miR-181c interaction has also been investigated in both mice and humans and found involved in various diseases, implying an important role of the 3’ UTR [40,41].

### The 3’ UTR of ND5 mRNA

The ND5 mRNA carries a 3’ UTR of ∼600 nt, a feature conserved among all the investigated species and developmental stages. Vertebrate ND5 mRNA appears unusual in several aspects. The ND5 mRNA stability was reported to be tightly regulated in mice and suggested to play a key role in respiratory control [42,43]. The ND5 mRNA appears to be the less structured among all human mitochondrial mRNAs [8], and we report this mRNA to be frequently targeted by m^6^A modifications, which is in line with the above observations since m^6^A is linked to mRNA stability [35]. The long ND5 mRNA 3’ UTR is antisense to the ND6 mRNA and tRNA-Glu. The non-polyadenylated ND6 mRNA has a heterogenic 3’ UTR of several hundred nucleotides, and is complementary to the coding part of ND5 mRNA [3,44]. Thus, the duplex region could be as long as 1 kb. The ND5 mRNA/ ND6 mRNA interaction complex may be a candidate in coordinating the H-strand and L-strand transcripts by antisense base-pairing. An interesting observation is that ND6 mRNA, but not ND5 mRNA, is closely associated with mtDNA at the inner mitochondrial membrane surface [45], suggesting that compartmentalization may take part in the regulation. Finally, our results from scissortail *Rasbora* indicates that lncND6 [3] is generated from the full-length ND5 mRNA by RNA cleavage only eight nucleotides downstream the UAA stop codon (S1 Table and S2A Figure).

### The mitochondrial long noncoding RNA lncOriL

The ∼300 nt lncOriL is polyadenylated (poly-A tail of 40-50 adenosines), highly structured, and shows a 70% sequence similarity of between zebrafish and human (Figure 5). We may only speculate why this ubiquitously expressed and relatively abundant mt-lncRNA has not been characterized previously. One plausible explanation is due to technical restrictions in the sequencing methodology. The fact that dRNA-seq, and not cDNA-based sequencing, detects lncOriL is favouring first-strand RT synthesis as a bottleneck. This notion is supported by ONT sequencing experiments in scissortail *Rasbora* using the same RNA sample in both approaches. While dRNA-seq readily detects lncOriL (S2A Figure), cDNA-based sequencing did not [10]. Interestingly, in a report by Saccone and co-workers published years ago, short H-strand specific RNA transcripts that spanned the OriL region were detected in rat liver mitochondria [46]. These analyses did not rely on first-strand RT synthesis, but on RNA protection and Northern hybridization. While the RNA candidates were not end-mapped or sequenced, it may still be plausible that a rat lncOriL homolog was among the detected fragments.

The fact that lncOriL is polyadenylated and ubiquitously expressed in teleost fish and mammalian mitochondria favour a biological role. lncOriL has a potential of being regulated by antisense RNA since the tRNA-derived fragment tiRNA5-Asn may form a strong duplex with lncOriL. tiRNA5-Asn was co-expressed with lncOriL in all zebrafish developmental stages and one of the most common tRNA-derived fragments in human tissue. How lncOriL eventually works in cells is currently not known, but one possibility could simply be by base-pairing to the L-strand mtDNA and thus preventing the formation of the functional essential OriL hairpin. Alternatively, lncOriL may function as a molecular sponge that sequester specific proteins important for mitochondrial import or function. The fact that the 11-bp stem in the OriL hairpin, that specifically binds mitochondrial DNA polymerase gamma [47], is identical in sequence to that of lncOriL, makes the DNA polymerase an interesting candidate. These, and other possibilities need to be further explored in experimental settings.

## Conclusions

In this study we characterized the poly-A mito-transcriptome in zebrafish during early development and compared our findings to other teleost fish and mammals, including human. We identified three unusual polyadenylated transcripts that carry significant regions of non-coding RNA sequences. The COI and ND5 mRNAs, which represent the most and less stable of the mRNAs, have 3’ untranslated regions containing antisense genes with regulatory potential. We described lncOriL, a previously undetected mitochondrial long-noncoding RNA that is polyadenylated and abundant in zebrafish early development, and present in all investigated teleost fish and mammals. lncOriL appears highly structured and covers the origin of light strand replication, and has the potential of being regulated in mitochondria by a tRNA-derived fragment in both zebrafish and human.

## Data availability statement

The raw sequencing m^6^A datasets have been deposited in the NCBI Sequence Read Archive under BioProject PRJNA1364744 (SRA accessions SRR36031735-SRR36031764). All raw sequencing data used for the mitochondrial tRNA-derived fragment profiling are publicly available from the NCBI Sequence Read Archive under BioProject accession PRJNA667163 (Saarland cohort). The human mitochondrial genome reference sequence (NC_012920.1) was obtained from NCBI GenBank. All Nanopore dRNA-seq datasets have been deposited in the NCBI Sequence Read Archive under BioProject PRJNA1432743 (SRA accession SRR37486486-SRR37486492, SRR37498323, SRR37499850, SRR37531269 and SRR37532071-SRR37532074.)

## Ethics statement

Experiments including zebrafish: The experimental process and husbandry were performed in agreement with the Norwegian Regulation experimental process and husbandry were performed in agreement with the Norwegian Regulation on Animal Experimentation (The Norwegian Animal Protection Act, No. 73 of 20 December 1974). This was certified by the National Animal Research Authority, Norway, General License for Fish Maintenance and Breeding no. 17. Experiments including human blood samples: The study was approved by the Regional Ethics Committee (REK Nord, approval numbers P REK Nord 32/2004 and 2013/1801/REK Nord) and the Norwegian Directorate of Health.

## Author contributions

TEJ: Study concept, Study design, Methodology, RNA sequencing handling, Nanopore sequencing, Nanopore sequencing data analysis, Writing – review & editing, Visualization, Supervision, Project administration.

AW: Study design, Methodology, Zebrafish developmental biology, m^6^A modification, RNA sequencing handling, Illumina sequencing, Illumina sequencing data analysis, Writing – review & editing.

SWP: Study design, Methodology, Scissortail samples, RNA sequencing handling, Sanger sequencing, Sanger sequencing data analysis, Writing – review & editing, Visualization.

CA: Methodology, Zebrafish husbandry management, Writing – review & editing. ÅEE: Methodology, Pig samples, Writing – review & editing.

ISJ: Methodology, Human samples, Writing – review & editing.

OLB: Methodology, Human samples, Writing – review & editing, Funding acquisition.

BOK: Study design, Methodology, Human samples, RNA sequencing handling, Nanopore sequencing data analysis, Writing – review & editing, Funding acquisition.

IB: Study concept, Study design, Methodology, Zebrafish developmental biology, m^6^A modification, Writing – review & editing, Visualization, Supervision, Project administration, Funding acquisition. SDJ: Study concept, Study design, Methodology, Nanopore sequencing data analysis, Writing – original draft, Writing – review & editing, Visualization, Supervision, Project administration, Funding acquisition.

## Acknowledgements

We thank Dr. Yong Peng, Dr Ruiqi Ge, and Dr Chuan He (University of Chicago) for support in m^6^A sequencing analyses. We also thank Dr. Lars Martin Jakt (Nord University) for support in R analyses and visualization. Biopsies from porcine was kindly provided by the Hemostatic emergency surgery course managed through the University Hospital of North Norway (UNN) and Nord University.

## Generative AI statement

The authors declare that no Generative AI was used in the creation of this manuscript.

## Supporting information

**S1 Figure.**
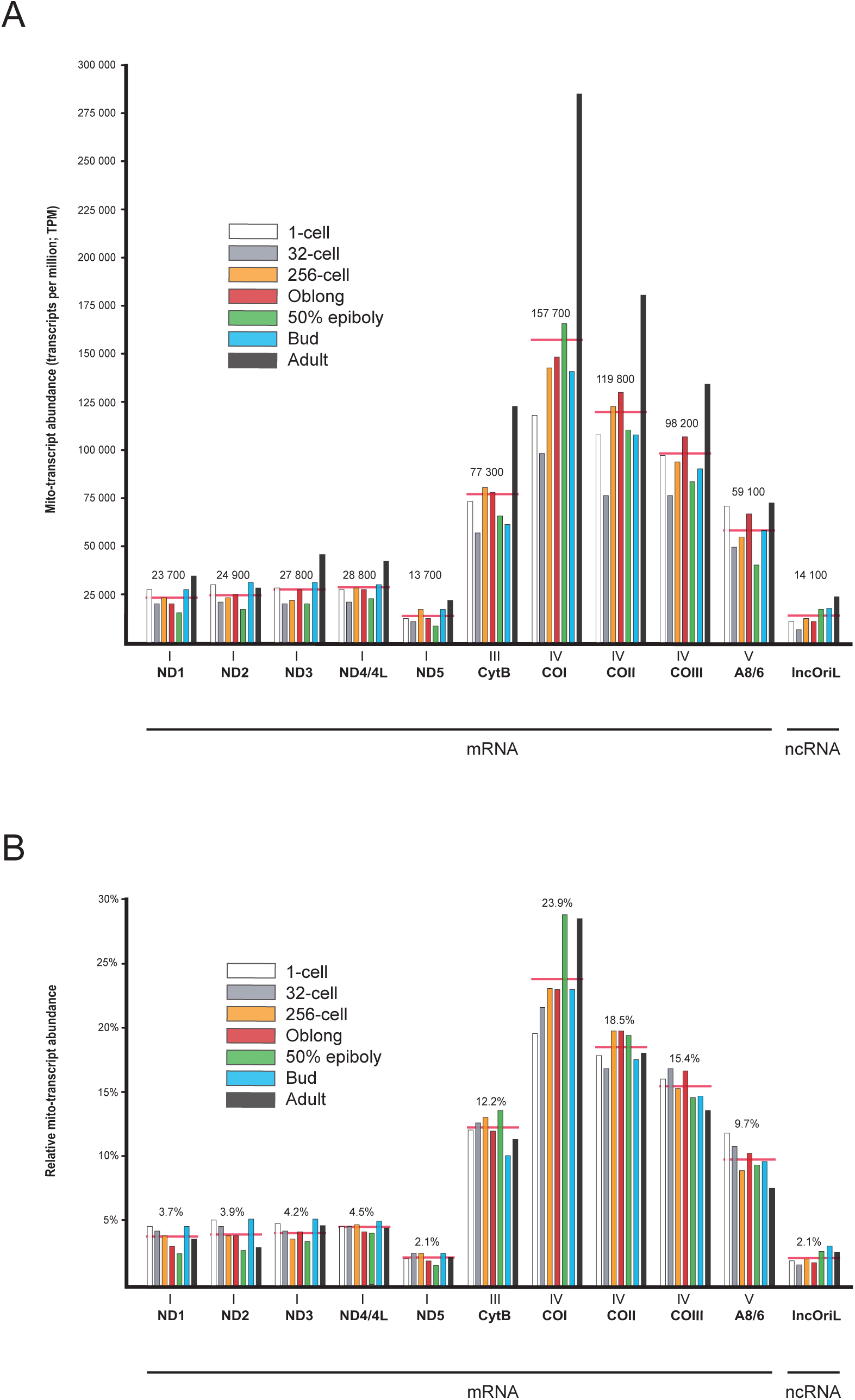
Enrichment of H-strand specific RNA during zebrafish early development. The enrichment level is indicated by **A**) transcripts per million (TPM), and **B**) Relative level (%) of TPM within each sample. The relative levels of individual mRNAs between stages were practically invariant, but different from the adult stage (apparent increase of Complex III and IV). lncOriL and ND5 mRNA (Complex I) have similar enrichment level.

**S2 Figure.**
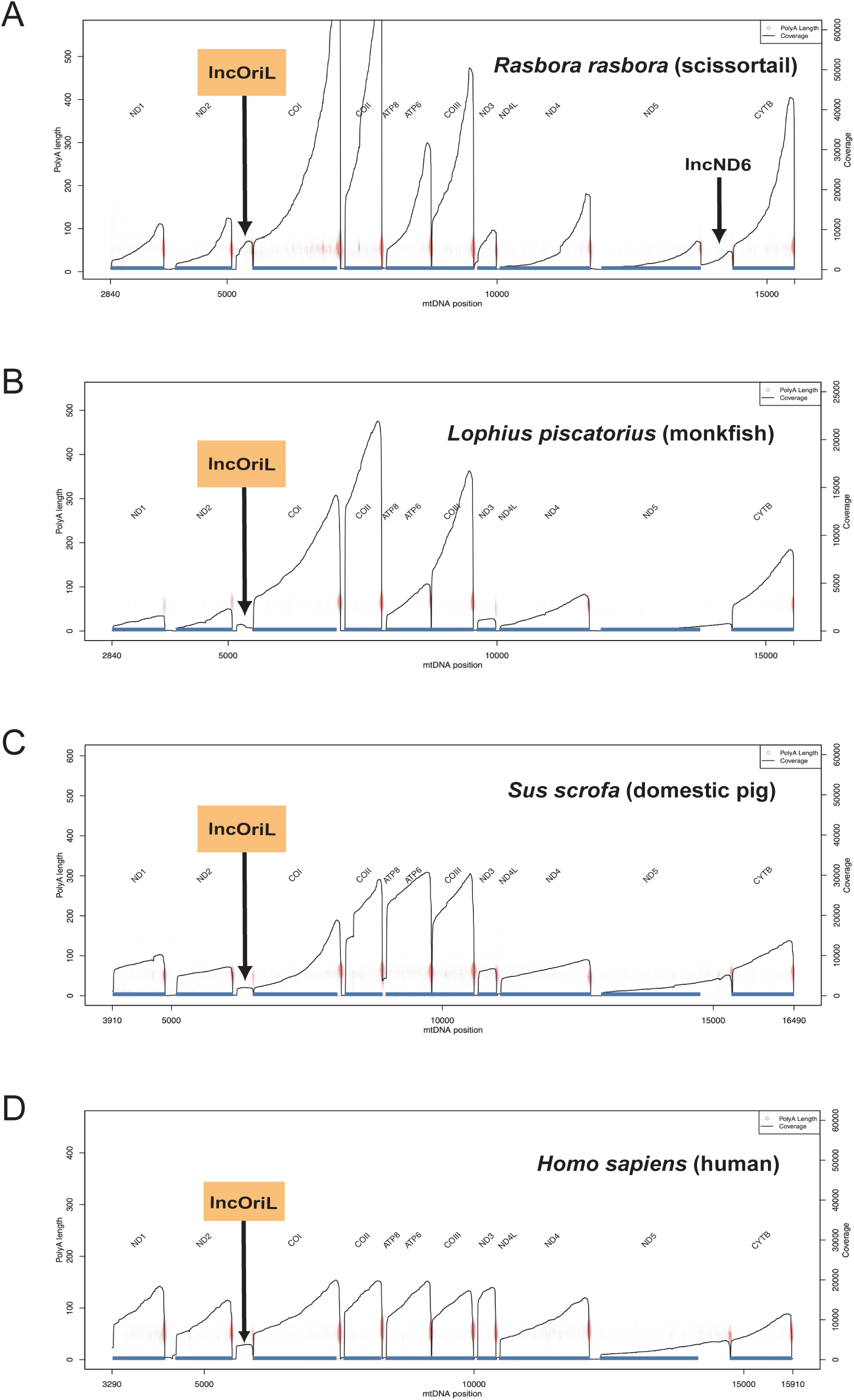
Mitochondrial transcriptome profiles of teleost fish and mammals, based on ONT dRNA-seq. IncOriL is indicated. **A**) *Rasbora rasbora* (scissortail) profile. lncND6 is indicated. **B**) *Lophius piscatorius* (monkfish) profile. **C**) *Sus scrofa* (pig) profile. **D**) *Homo sapiens* (human) profile.

**S3 Figure.**
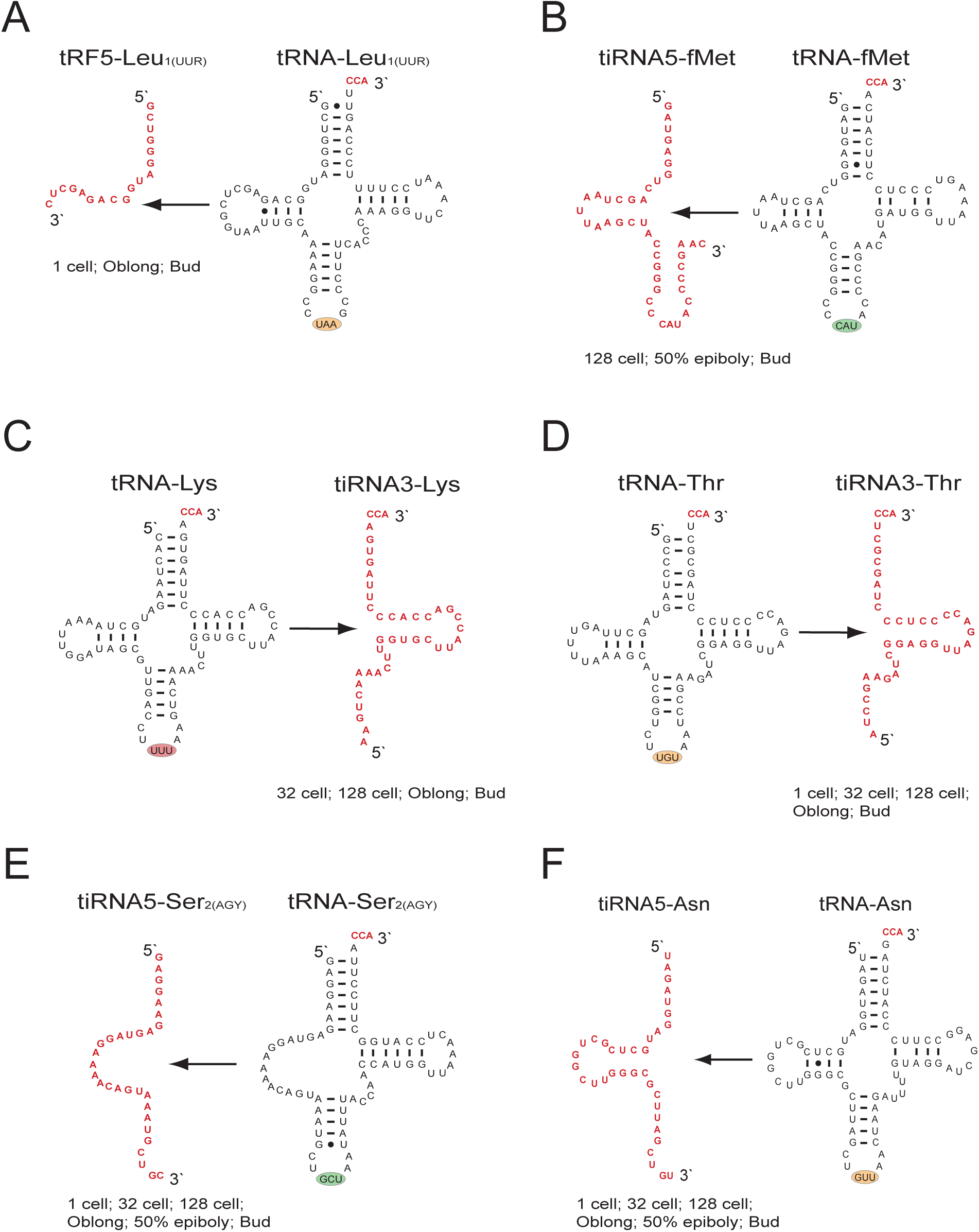
tRNA-derived small RNAs generated during zebrafish early development. **A**) tRF5-Leu_1(UUR)_; **B**) tiRNA5-fMet; **C**) tiRNA5-Lys; **D**) tiRNA3-Thr; **E**) tiRNA5-Ser_2(AGY)_; **F**) tiRNA3-Asn. All 3’ fragments contain the terminal CCA motif, which indicate a mature tRNA origin.

**Table S1.**
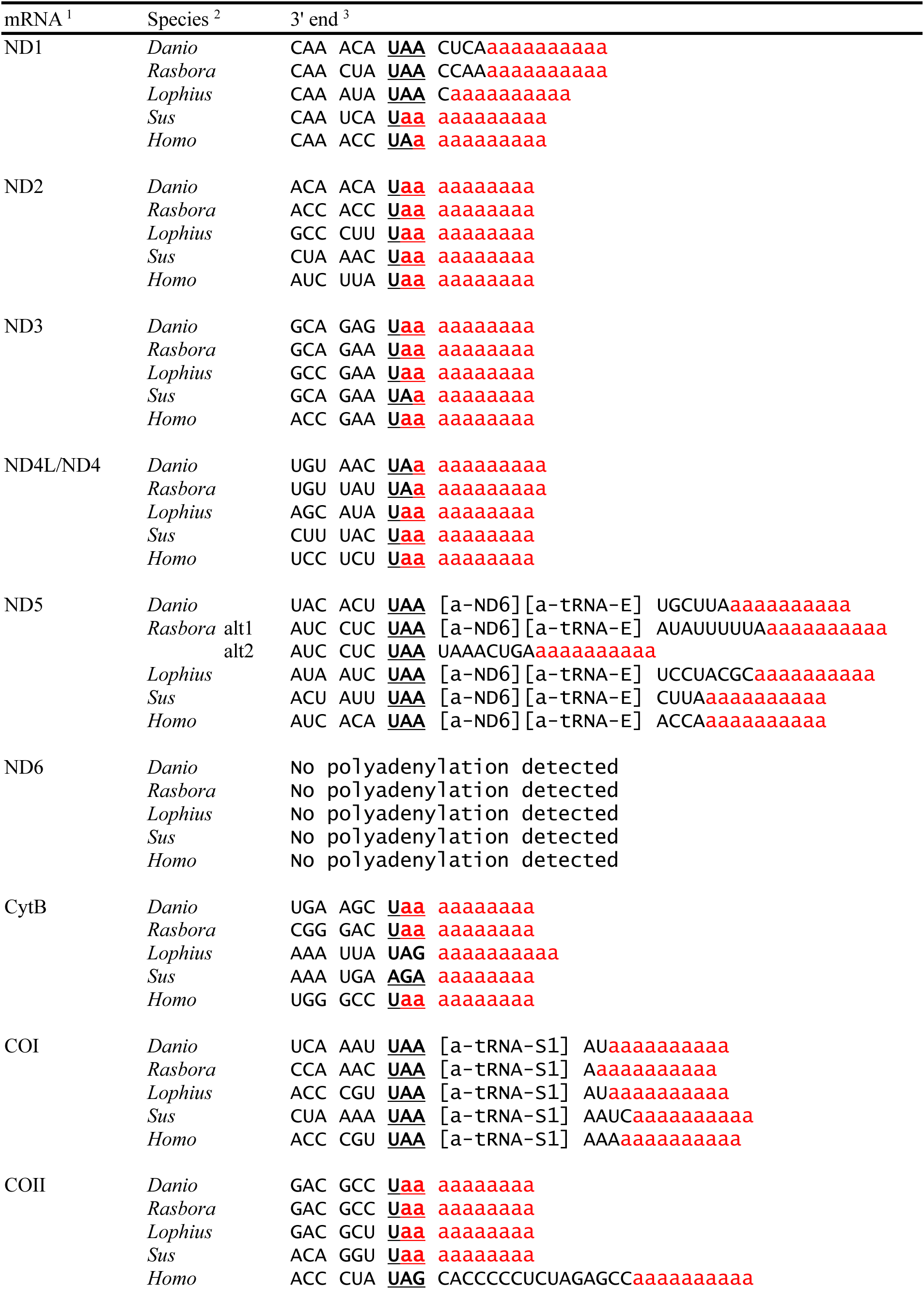
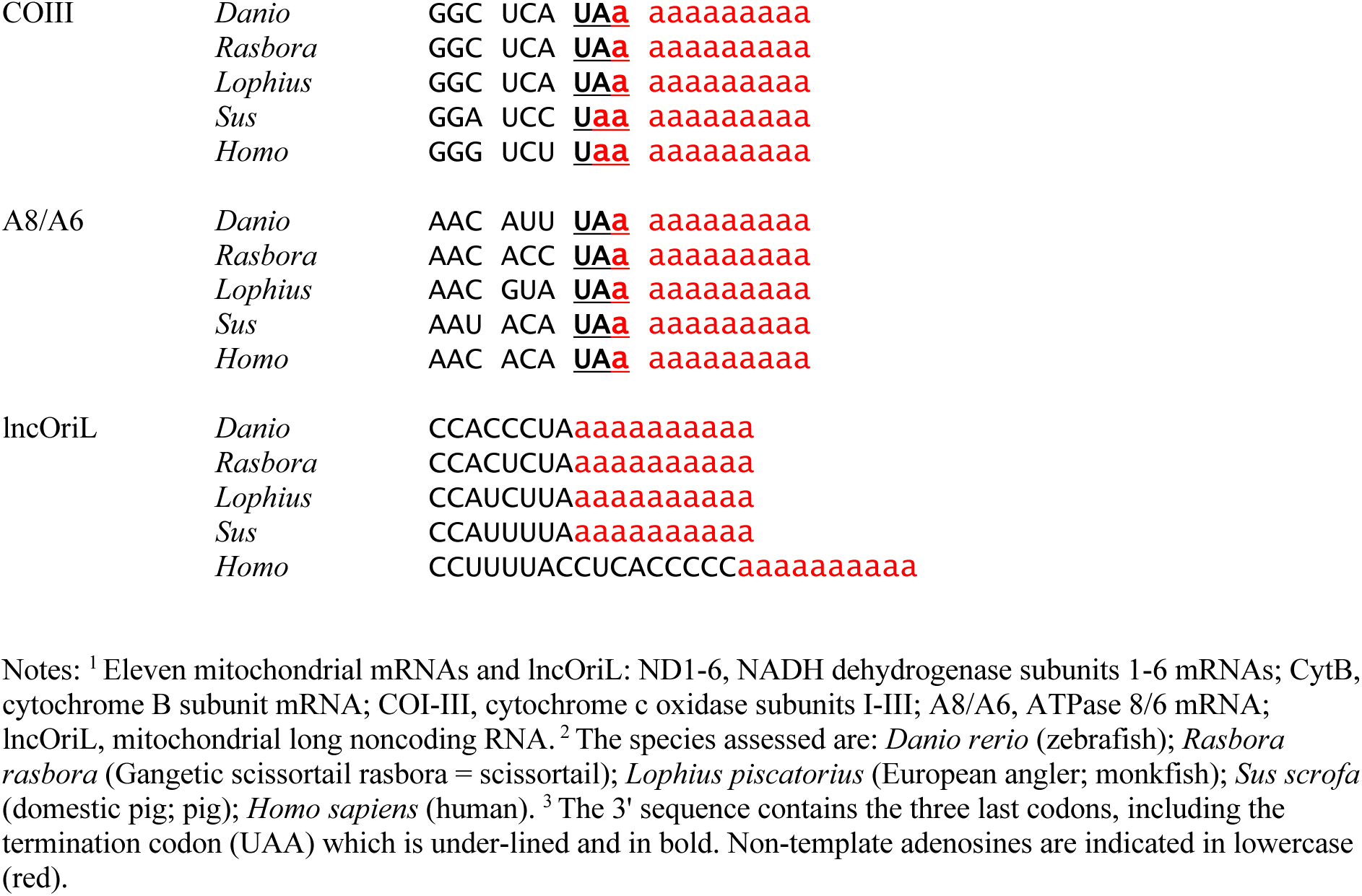
Poly-A site features in mitochondrial mRNAs and lncOriL assessed in zebrafish, scissortail, monkfish, pig, and human. Data obtained from ONT direct RNA sequencing.

**Table S2.** Mapped m^6^A sites in zebrafish mitochondrial RNAs (DRACH motifs) during early development.

| m6A site<sup>1</sup> | Motif (DRACH) | RNA<sup>2</sup> | 1-cell | 256-cell | Oblong | 50% Epi | Bud |
| --- | --- | --- | --- | --- | --- | --- | --- |
| A3524-H | GGACC | ND1 | - | - | - | - | + |
| A3827-H | UAA<u>CU</u> | ND1 (term) | - | - | - | + | + |
| T4299-L | UAA<u>CA</u> | a-ND2 | - | - | - | - | + |
| A4418-H | UG<u>ACA</u> | ND2 | - | + | - | - | - |
| A4423-H | GG<u>ACU</u> | ND2 | + | - | - | - | - |
| A4744-H | UG<u>ACC</u> | ND2 | + | + | - | - | + |
| A5047-H | GG<u>ACU</u> | ND2 | - | - | + | - | - |
| A5060-H | UG<u>ACC</u> | ND2 | - | + | - | - | - |
| A7514-H | AG<u>ACU</u> | COII | - | - | - | - | + |
| A8050-H | UG<u>ACC</u> | ATPase 8 | + | - | - | - | - |
| A8431-H | UG<u>ACU</u> | ATPase 6 | + | + | - | + | - |
| A8482-H | GG<u>ACA</u> | ATPase 6 | - | + | - | - | - |
| A8832-H | UG<u>ACC</u> | COIII | - | - | - | + | - |
| A8905-H | UG<u>ACA</u> | COIII | - | - | - | + | - |
| A9158-H | UG<u>ACC</u> | COIII | - | - | - | - | + |
| T9399-L | AG<u>ACC</u> | a-COIII | - | + | - | - | - |
| T12087-L | AG<u>ACU</u> | a-ND5 | + | - | - | - | - |
| A12125-H | AG<u>ACC</u> | ND5 | + | + | - | - | - |
| A12577-H | UG<u>ACC</u> | ND5 | + | + | - | - | - |
| A13013-H | AG<u>ACA</u> | ND5 | +(100%) | - | - | - | +(100%) |
| A13127-H | AG<u>ACG</u> | ND5 | + | - | - | - | - |
| A13273-H | AG<u>ACA</u> | ND5 | +(100%) | - | +(100%) | - | - |
| A13481-H | UAA<u>CA</u> | ND5 | - | + | - | - | - |
| A13629-H | GG<u>ACC</u> | ND5 | + | - | - | - | - |
| A13679-H | GG<u>ACC</u> | ND5 | - | + | - | - | - |
| A13794-H | AG<u>ACC</u> | a-ND6 | - | - | + | - | - |
| T14477-L | AG<u>ACA</u> | a-CytB | +(100%) | - | - | +(100%) | - |
| A15040-H | AG<u>ACC</u> | CytB | - | - | - | - | + |

Notes: <sup>1</sup> Eleven mitochondrial mRNAs and lncOriL: ND1-6, NADH dehydrogenase subunits 1-6 mRNAs; CytB, cytochrome B subunit mRNA; COI-III, cytochrome c oxidase subunits I-III; A8/A6, ATPase 8/6 mRNA; lncOriL, mitochondrial long noncoding RNA. <sup>2</sup> The species assessed are: *Danio rerio* (zebrafish); *Rasbora rasbora* (Gangetic scissortail rasbora; scissortail); *Lophius piscatorius* (European angler; monkfish); *Sus scrofa* (domestic pig; pig); *Homo sapiens* (human). <sup>3</sup> The 3' sequence contains the three last codons, including the termination codon (UAA) which is under-lined and in bold. Non-template adenosines are indicated in lowercase (red).

**Table S3.** Poly-A tail length in zebrafish, scissortail, monkfish, pig, and human based on ONT direct RNA sequencing. “All” (dark gray boxes), refers to average of all 10 mRNAs and lncOriL together. lncOriL values are indicated by light gray boxes.

| Specie | mRNA | Mean | Median |
| --- | --- | --- | --- |
| *Danio rerio* (zebrafish) | All – 1-cell | 62 A | 64 A |
|  | All – 32-cell | 60 A | 62 A |
|  | All – 256-cell | 57 A | 59 A |
|  | All – 50% epiboly | 58 A | 59 A |
|  | All – oblong | 62 A | 63 A |
|  | All – bud | 61 A | 61 A |
|  | All – adult | 55 A | 56 A |
|  | COI – 1-cell | 54 A | 56 A |
|  | COI – 32-cell | 54 A | 55 A |
|  | COI – 256-cell | 51 A | 53 A |
|  | COI – 50% epiboly | 53 A | 54 A |
|  | COI – oblong | 55 A | 56 A |
|  | COI – bud | 54 A | 55 A |
|  | COI – adult | 50 A | 51 A |
|  | ND5 – 1-cell | 54 A | 56 A |
|  | ND5 – 32-cell | 53 A | 56 A |
|  | ND5 – 256-cell | 49 A | 52 A |
|  | ND5 – 50% epiboly | 50 A | 52 A |
|  | ND5 – oblong | 56 A | 58 A |
|  | ND5 – bud | 56 A | 58 A |
|  | ND5 – adult | 51 A | 52 A |
|  | lncOriL – 1-cell | 41 A | 38 A |
|  | lncOriL – 32-cell | 40 A | 38 A |
|  | lncOriL – 256-cell | 41 A | 40 A |
|  | lncOriL – 50% epiboly | 46 A | 46 A |
|  | lncOriL – oblong | 48 A | 47 A |
|  | lncOriL – bud | 49 A | 49 A |
|  | lncOriL – adult | 50 A | 50 A |
| *Rasbora rasbora* (scissortail) | All – adult | 54 A | 54 A |
|  | COI – adult | 53 A | 52 A |
|  | ND5 – adult | 46 A | 48 A |
|  | lncOriL – adult | 48 A | 48 A |
| *Lophius piscatorius* (monkfish) | All – adult | 64 A | 64 A |
|  | COI – adult | 63 A | 65 A |
|  | ND5 – adult | 43 A | 42 A |
|  | lncOriL – adult | 41 A | 40 A |
| *Sus scrofa* (pig) | All – adult | 60 A | 59 A |
|  | COI – adult | 60 A | 61 A |
|  | ND5 – adult | 52 A | 51 A |
|  | lncOriL – adult | 51 A | 50 A |
| *Homo sapiens* (human) | All – Ex1, T=0 | 53 A | 53 A |
|  | All – Ex1, T=120 | 54 A | 54 A |
|  | All – Ex2, T=0 | 51 A | 52 A |
|  | All – Ex2, T=120 | 52 A | 53 A |
|  | COI – Ex1, T=0 | 52 A | 53 A |
|  | ND5 – Ex1, T=0 | 46 A | 47 A |
|  | lncOriL – Ex1, T=0 | 44 A | 44 A |

